# Dissecting the nanoscale lipid profile of caveolae

**DOI:** 10.1101/2020.01.16.909408

**Authors:** Yong Zhou, Nicholas Ariotti, James Rae, Hong Liang, Vikas Tillu, Shern Tee, Michele Bastiani, Adekunle T. Bademosi, Brett M. Collins, Frederic A. Meunier, John F. Hancock, Robert G. Parton

## Abstract

Caveolae are specialized domains of the vertebrate cell surface with a well-defined morphology and crucial roles in cell migration and mechanoprotection. Unique compositions of proteins and lipids determine membrane architectures. The precise caveolar lipid profile and the roles of the major caveolar structural proteins, caveolins and cavins, in selectively sorting lipids have not been defined. Here we used quantitative nanoscale lipid mapping together with molecular dynamic simulations to define the caveolar lipid profile. We show that caveolin1 (CAV1) and cavin1 individually sort distinct plasma membrane lipids. Intact caveolar structures composed of both CAV1 and cavin1 further generate a unique lipid nano-environment. The caveolar lipid sorting capability includes selectivities for lipid headgroups and acyl chains. Because lipid headgroup metabolism and acyl chain remodelling are tightly regulated, this selective lipid sorting may allow caveolae to act as transit hubs to direct communications among lipid metabolism, vesicular trafficking and signalling.

---

Caveolae are a striking morphological feature of the plasma membrane of many vertebrate cells. Caveolae have been implicated in mechanoprotection, endocytosis, signal transduction and lipid regulation ^1-6^. The characteristic morphology of caveolae, with a bulb connected to the plasma membrane by a highly-curved neck, is generated by integral membrane proteins termed caveolins and by lipid binding peripheral membrane proteins, the cavins ^7-14^. Specifically, caveolin-1 (CAV1) and caveolin-3 (CAV3, in striated muscle) and cavin1 (or polymerase I and transcript release factor, PTRF) are essential for caveola formation ^7,10^. Another set of key caveolar components comprise the plasma membrane (PM) lipids. While biophysical studies have consistently suggested that the lateral distribution of lipids determines and/or responds to changing membrane morphology, our understanding of the lipid composition of caveolae and how this contributes to caveola formation, function, and disassembly, is still relatively primitive. A detailed molecular understanding of the lipid constituents of caveolae, defined both by their headgroups and acyl chains, is crucial for understanding the formation of caveolae, as well as their disassembly, processes that are crucial for caveola function. Moreover, caveolae can provide a paradigm for understanding how local concentrations of specific lipid species contribute to membrane morphogenesis. Further, many lipid messengers directly participate in cell surface signalling. Thus, a better understanding of the caveolar lipid profile will shed more light on the biological significance of caveolae. In this study, we combined model cellular systems with lipid depletion/rescue experiments and a quantitative ultrastructural mapping technique that has been used to define the nanoscale lipid association of Ras isoforms ^15-20^.

## A model system for *de novo* assembly of caveolae by caveolin and cavin1

To dissect the selective lipid sorting of each caveolar associated protein we used MCF7 cells which lack expression of both caveolins and cavins ^21 22^. We first expressed the basic components of caveolae, CAV1 and cavin1 (Figure 1A), to assemble caveolae *de novo*. Expression of CAV1 and cavin1 in MCF7 cells was sufficient to generate surface pits with the typical features of caveolae as judged by immunoelectron microscopy on frozen sections (Figure 1B), by conventional EM with a surface stain (Figure 1C), and by using a nanobody fused to the ascorbate peroxidase, APEX2, to detect YFP-CAV1 (Figure S1). PM sheets labeled for the expressed proteins showed co-association of CAV1 and cavin1 with structures of caveolar morphology (Figure 1D) whereas Cavin1 expressed alone showed a similar level of surface labeling (Figure S2A) but no detectable association with defined membrane domains (Figure 1E). In contrast to CAV1 expressed with cavin1, CAV1 expressed alone showed a heterogeneous pattern of PM labeling when localized using the nanobody-APEX system (Figure S1).

**Figure 1.**
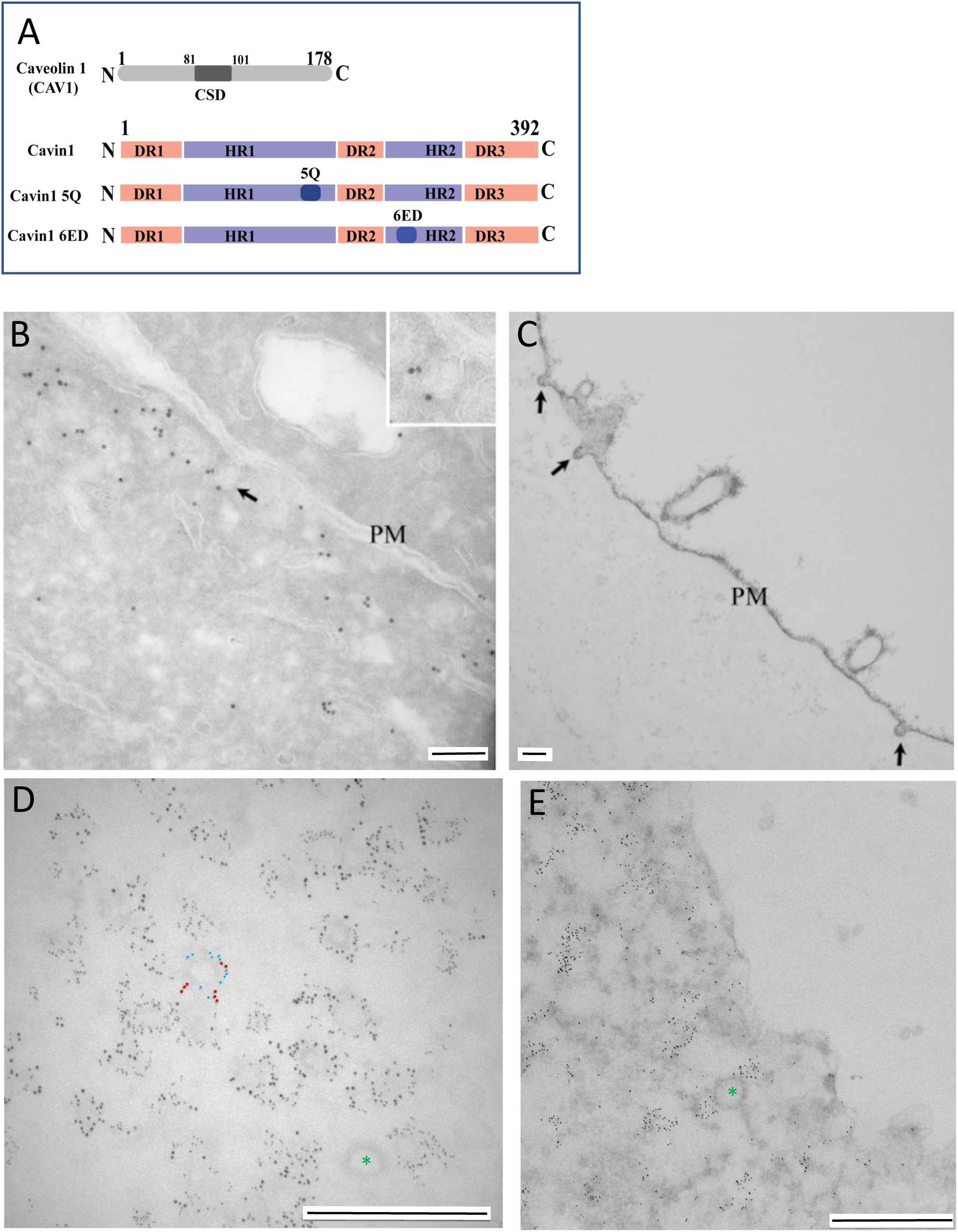
Expression of CAV1, cavin1, and mutants in MCF7 cells. A) Domain structure of caveolin (CAV1), wild type cavin1, and cavin1 mutants used in this study. CSD; caveolin scaffolding domain; HR, helical region; DR, disordered region. Cavin1 5Q and Cavin1 6ED denote amino acid substitutions in the putative PI(4,5)P_2_–binding region of HR1 and in the UC1 domain of HR2, respectively. B-E. MCF7 cells expressing CAV1 and cavin1 (B, C, D) or cavin1 alone (E); B. Frozen section showing labeling for CAV1 in MCF7 cells co-expressing CAV1 and cavin1. Labeling is associated with membranous profiles close to the surface, some of which are connected to the plasma membrane (eg. see arrow; shown at higher magnification in the inset). C; ruthenium red labeled plasma membrane (PM) showing caveolae (arrows). D; sonicated plasma membrane sheet showing labeling for CAV1-mCherry (2nm gold) and Cavin1-GFP (6nm gold) associated with approx. 60-80nm diameter profiles characteristic of caveolae. In panel B the large gold (red; cavin1) and the small gold (blue; CAV1) have been highlighted on one structure (raw image Figure S2). E; sonicated plasma membrane sheet showing clustered labeling for Cavin1-GFP (6nm gold). Clathrin coated pits (green asterisks) are unlabeled. Bars; C,D 200nm; D,E 500nm.

We further characterized the features of this putative non-caveolar CAV1 pool. Single molecule tracking of CAV1 tagged with monomeric Eos2 showed that CAV1 expressed alone in MCF7 cells showed greater mobility than in MDCK cells that have endogenous caveolae. Analysis of the mean square displacement (MSD) of CAV1-mEos2 mobility revealed an increase in confinement in MDCK cells as shown by the MSD curves and the area under the MSD curves (Figure S1C). These results are consistent with the reported effect of cavin1 KD on CAV1 diffusion as shown by FRAP ^7^ and super-resolution light microscopy ^23^. APEX staining for YFP-CAV1 was localized to discrete patches of the PM, in areas enriched in vesicles and tubular profiles (Figure S1D, E) and was also associated with pits and vesicles of varying diameter (eg. Figure S1F). This pattern of staining was quite distinct from the restriction of APEX labeling to structures with caveolar morphology in cells co-transfected with cavin1 (Figure S1G).

## Caveolae show a distinct lipid profile

Having established a system in which we can assess the surface distribution of individually expressed CAV1 and cavin1, as well as the two proteins co-expressed (caveolae), we then used immunogold EM spatial mapping to quantify the nanoscale organization of the CAV1 and cavin1 domains. This system allows an unbiased quantitative analysis of the clustering of a particular type of proteins (univariate analysis) or their association with other proteins (bivariate analysis).

For assessment of caveolar localization in cells expressing both CAV1 and cavin1, labeling of CAV1 was used as the caveolar marker. CAV1, expressed alone or with cavin1 in MCF7 cells, labeled domains of approximately 40nm radius (Figure S2B). The lateral univariate clustering of the gold labelling within a select 1 μm^2^ PM area was calculated using the Ripley’s univariate K-function analysis. The extent of nanoclustering, *L*(*r*)-*r*, was plotted as a function of the cluster radius, *r*, in nanometers (Figure S2B). The *L*(*r*)-*r* value of 1 indicates the 99% confidence interval (C.I.), the values above which indicate statistically meaningful clustering. The peak *L*(*r*)-*r* value is termed as Lmax and summarizes nanoclustering statistics. A larger Lmax indicates more extensive nanoclustering. The spatial analysis showed a more restricted cluster size for CAV1 when expressed with cavin1 than when expressed alone (Figure S2B). We next sought to determine the lipids co-localizing with the singly expressed proteins and the CAV1/cavin1 complex. PM sheets were prepared from MCF7 cells co-expressing RFP-labeled caveolar proteins together with a lipid-binding domain: GFP-Lact-C2, GFP-PH-PLCδ, GFP-Spo20, GFP-PH-AKT, or GFP-D4H that specifically bind to PtdSer, PtdIns(4,5)P_2_, phosphatidic acid (PA), phosphatidylinositol 4,5-trisphosphate (PtdIns(3,4,5)P_3_) or cholesterol, respectively. PM sheets were co-labeled with anti-RFP-2nm-gold and anti-GFP-6nm-gold. The intact PM sheets were imaging using transmission EM at a magnification of 100,000X. The lateral co-localization of the two populations of gold particles within a select 1 μm^2^ PM area was calculated using the Ripley’s bivariate K-function. The extent of co-clustering, *L*_*biv*_(*r*)-*r*, was plotted as a function of the cluster radius, *r* (Figure 2A),. The *L*_*biv*_(*r*)-*r* value of 1 indicates the 95% C.I., the values above which indicate statistically meaningful co-localization. Area-under-the-curve between the *r* values of 10 and 110nm yields L-function-bivariate integrated (*<u>LBI</u>*), which summarizes the extent of co-localization. Larger LBI values indicate more extensive co-localization, with an LBI value of 100 as the 95% C.I.

**Figure 2.**
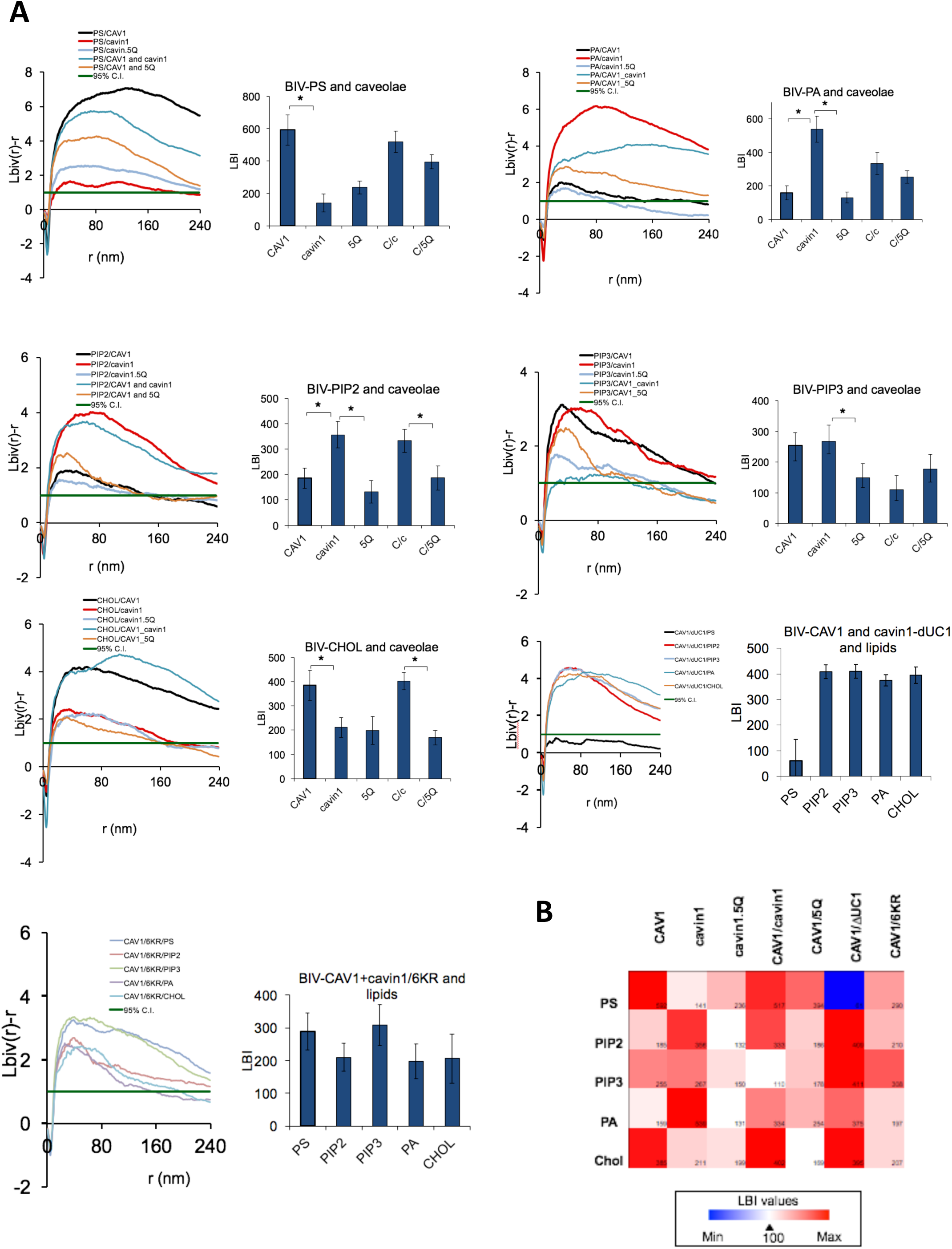
Lipid mapping of caveolar components expressed in a model caveola-deficient cell system. **A.** Bivariate clustering analyses of the indicated caveolar proteins and lipid-binding probes co-expressed in MCF7 cells; 5Q – Cavin1.5Q; C/c – CAV1/cavin1; C/5Q CAV1 plus cavin1.5Q; **B.** Bivariate clustering heatmap summary.

Singly expressed CAV1 statistically significantly associated with PtdSer, PtdIns(3,4,5)P_3_, and cholesterol (Figure 2A, B). In contrast, Cavin1 expressed alone preferentially co-clustered with PtdIns(4,5)P_2_, PtdIns(3,4,5)P_3_, and PA. This indicated that CAV1 and cavin1 sort different PM lipids. Next, CAV1 when expressed with cavin1 shows a quantitatively distinct lipid association profile to CAV1 expressed alone; association with PtdIns(3,4,5)P_3_ was decreased but association with PtdIns(4,5)P_2_ and PA was significantly increased (for summary schemes, see Figure 6). Taken together, CAV1 / cavin1 alone, or combined each sorts distinct set of lipids.

**Figure 3.**
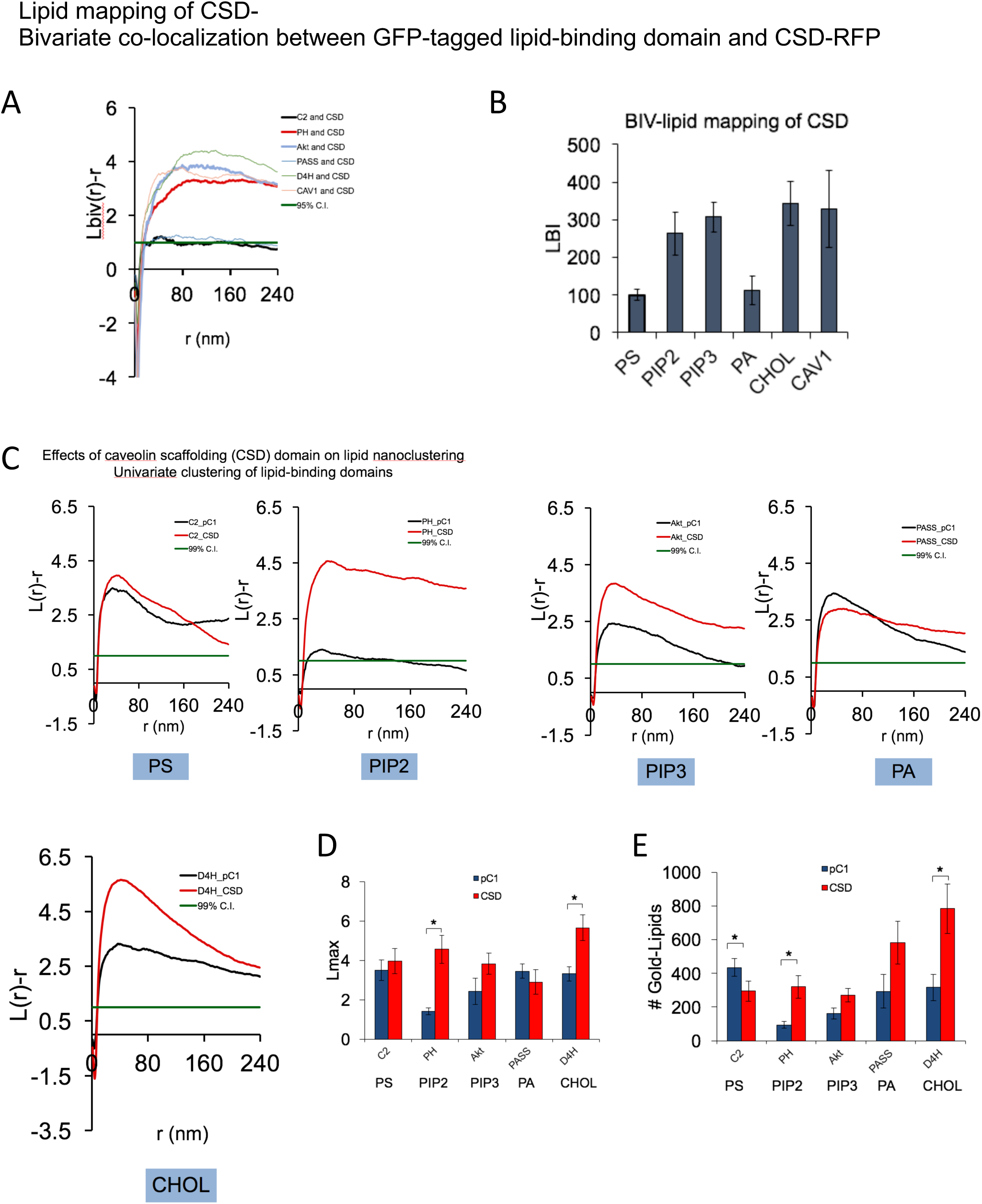
The caveolin scaffolding domain (CSD) shows a distinct lipid specificity, colocalizes with CAV1 and with eNOS and affects plasma membrane lipid organization. **A.** Bivariate analysis: CSD co-clusters with lipids and CAV1. **B.** Bivariate clustering summary. **C.** CSD affects PIP2, PIP3 and cholesterol nanoclustering. D. Lipid univariate clustering summary. **E.** CSD affects surface levels of PS and cholesterol.

**Figure 4.**
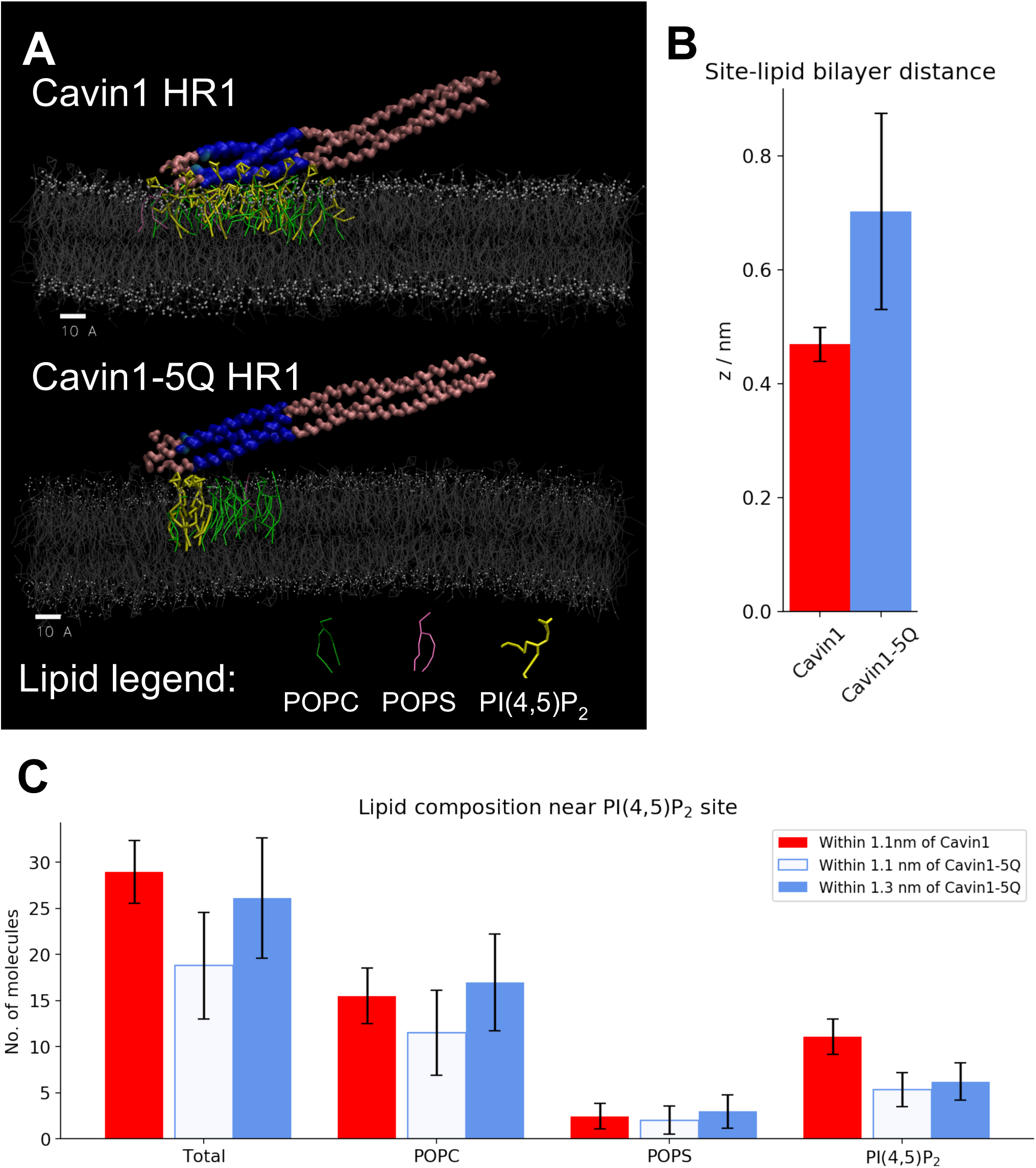
Coarse-grained molecular simulations of Cavin1 HR1 and Cavin1-5Q HR1 interacting with a model lipid bilayer. **A**. Side view of typical lipid bilayer-interacting configurations for (top) Cavin1 HR1 and (bottom) Cavin1-5Q HR1. Highlighted lipids are within 1.1nm of the Cavin1 interaction site (blue) or within 1.3nm of the Cavin1-5Q interaction site (blue). **B.** Minimum distance between the interaction site and lipid bilayer surface. **C.** Total number of lipids and number of lipids by type near the interaction site, with values for two different interaction distances shown for Cavin1-5Q.

**Figure 5.**
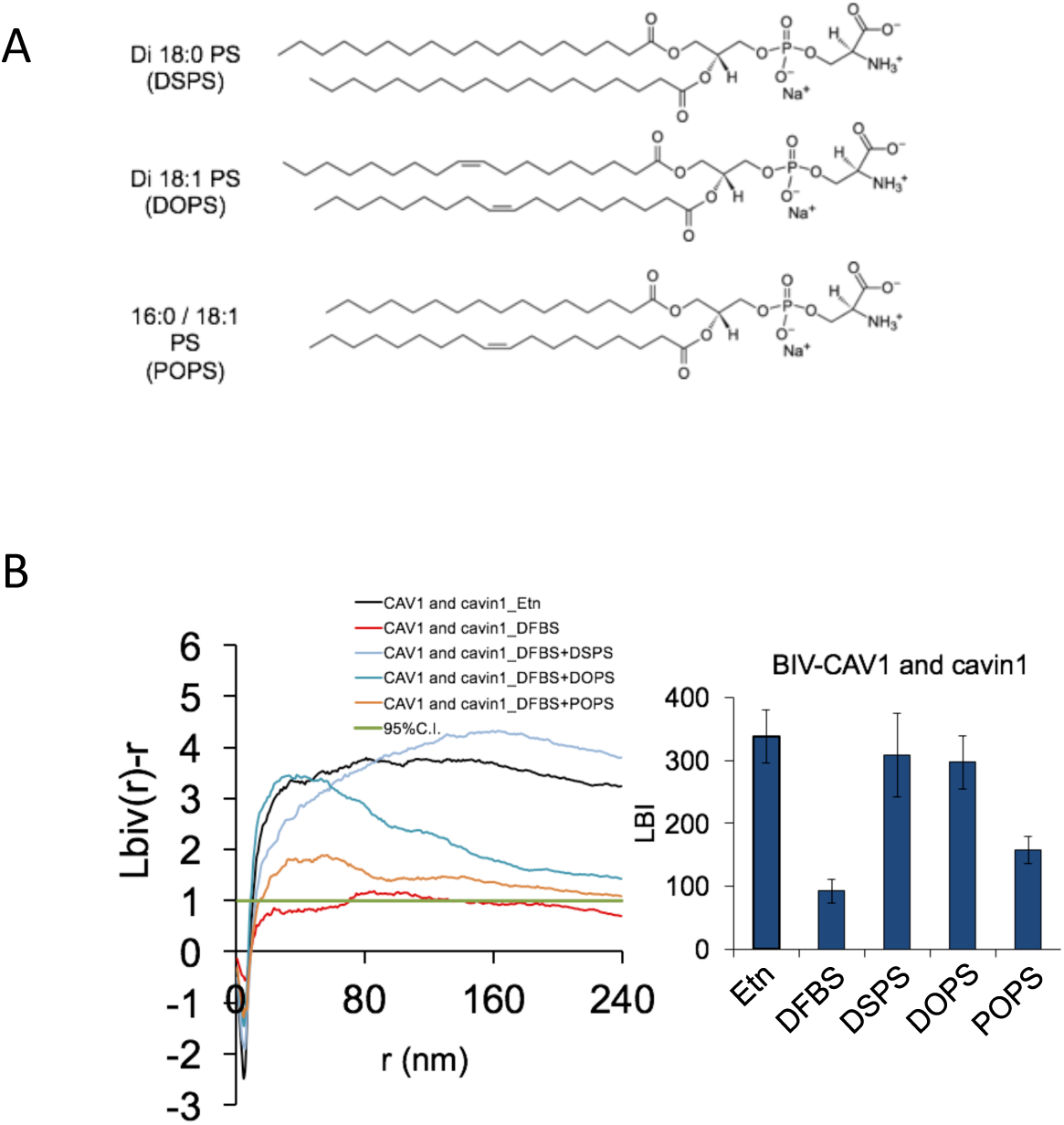
CAV1/cavin association is dependent on PS. **A.** Structure of different synthetic PS species used in this study. **B.** CAV1/cavin1 association is negligible in cells deficient in PS but restored in cells cultured in ethanolamine. Different PS species have distinct abilities to mediate CAV1/cavin1 association. Fully saturated DSPS and mono-unsaturated DOPS are effective at driving CAV1/cavin1 association but mixed chain POPS is ineffective.

**Figure 6.**
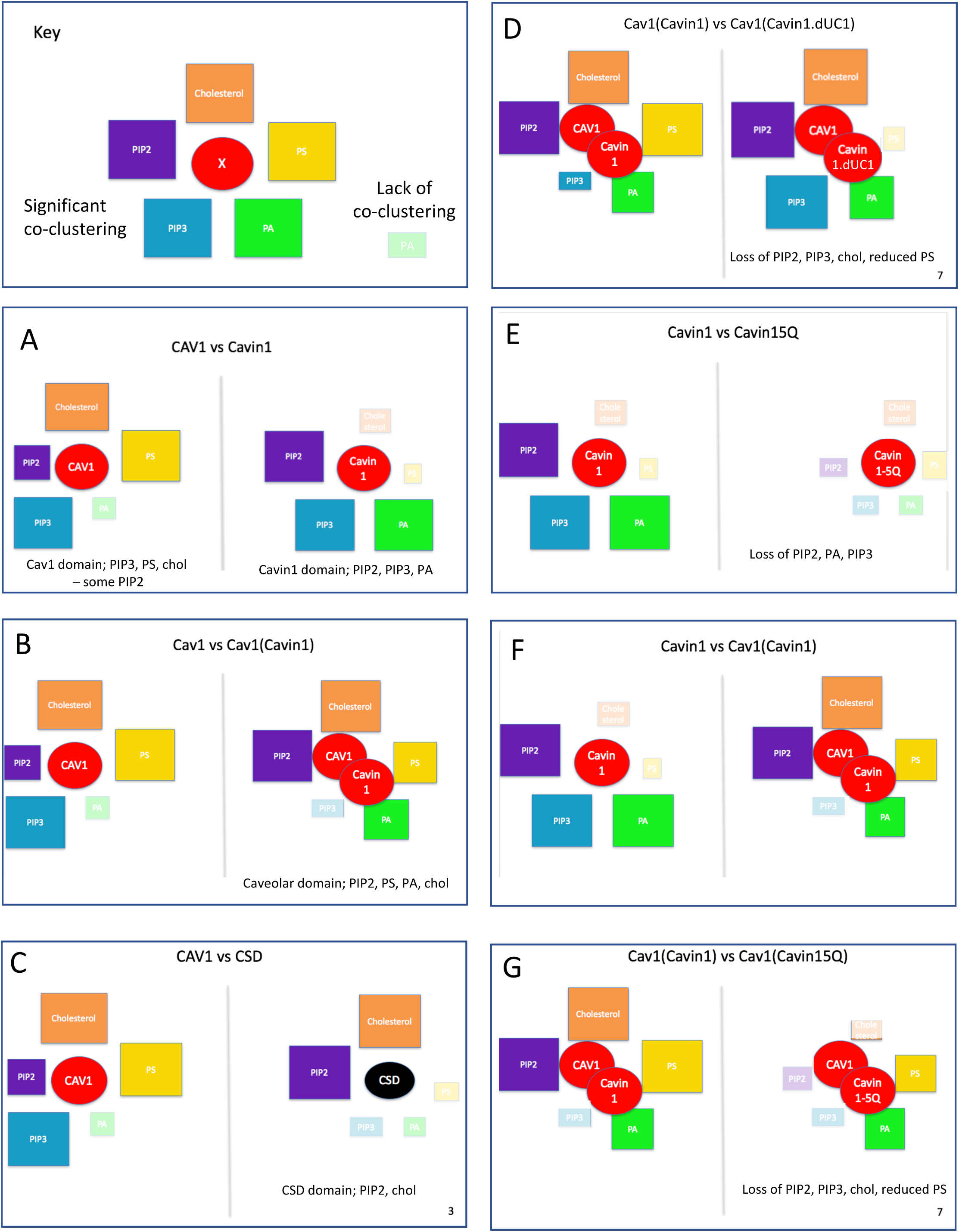
Schematic comparison of the lipid profiles associated with the indicated proteins or protein pairs (circles). Colors of square boxes indicate lipid species; the size of the box indicates the level of co-clustering with the indicated proteins (faded boxes indicate statistically insignificant association).

## The Caveolin scaffolding domain contributes to the selective lipid sorting of caveolae

To dissect the properties of CAV1 that could dictate association with specific lipid domains, we focussed on the conserved membrane proximal region of CAV1 termed the caveolin scaffolding domain, CSD (Figure 1A; Figure 3). The CSD is a potent and specific inhibitor of eNOS activity *in vitro* and *in vivo* ^24-27^ and has been reported to have membrane binding activity ^28,29^. We first examined the specific lipid profile (probed by the GFP-tagged lipid-binding domains) of the expressed RFP-CSD in an EM-bivariate co-localization analysis. The RFP-CSD showed significant PM labelling and co-clustered with PtdIns(4,5)P_2_, PtdIns(3,4,5)P_3,_ and cholesterol (Figure 3A,B). Unlike full length CAV1, however, the CSD alone showed no significant association with PtdSer (Figure 3B). In view of the association of the CSD with distinct lipid domains and the profound effects on signal transduction ^24,26^, we hypothesized that the CSD might affect nanoscale lipid organization. To test this, the RFP-tagged CSD was co-expressed with the suite of GFP-tagged lipid markers and their spatial distribution was quantified via the univariate spatial analysis. Expression of the CSD increased the clustering of PtdIns(4,5)P_2_ and cholesterol (Figure 3C, D). Counting the gold particles within the same 1 μm^2^ PM area estimates the surface density of the lipid probes. The presence of the CSD domain increased levels of PtdIns(4,5)P_2_, and cholesterol but caused a significant decrease in PtdSer (Figure 3E). These results indicate that the caveolin CSD co-localizes with a distinct set of PM lipids, and can have profound effects on surface spatial organization and levels of PM lipids in distinct manners.

## Cavin1 HR1 and UC1 domains contribute distinct lipid sorting capabilities

We next focussed on the other key structural protein, cavin1, and first investigated the role of the conserved PtdIns(4,5)P_2_ binding site in the N-terminal helical domain (HR1, residues 106-135) ^30^. Substitution of five key positively charged amino acids to glutamines within the HR1 domain (5Q; Figure 1A) disrupts binding of purified cavin1 to PtdIns(4,5)P_2_ in liposome binding assays. We therefore tested whether the co-clustering of specific lipids with cavin1 or with the CAV1/cavin1 complex would be affected by the 5Q substitutions in parallel bivariate lipid mapping amalysis. The 5Q substitutions in cavin1 caused a dramatic change in the association with PtdIns(4,5)P_2_, PtdIns(3,4,5)P_3_ and PA (Figure 2A,B). Co-expression of CAV1 and cavin1-5Q showed a decrease in association of PtdIns(4,5)P_2_, as well as a loss of cholesterol association, when compared to CAV1 expressed with wild type cavin1 (heatmap in Figure 2B). As shown in the heatmap in Figure 2B and the scheme in Figure 6, the net result of the loss of PtdIns binding through the 5Q site extended to the entire tested lipid network.

To further characterize the potential membrane lipid interactions of cavin1, we performed coarse-grained molecular dynamics (CG-MD) simulations using the MARTINI 2.2 force field ^31-33^. The cavin1 HR1 domain trimer and its 5Q mutant were modeled using PDB structure 4QKV ^30^ as the initial configuration, and the interaction of each protein with a bilayer composed of POPC, POPS and PI(4,5)P_2_ in an 80:15:5 ratio was observed over four runs of 3 μs each. In these simulations, the PI(4,5)P_2_-binding HR1 invariably approached the bilayer within hundreds of nanoseconds, even while the interaction between the rest of the protein and the bilayer was more variable, for both cavin1-HR1 and cavin1-5Q HR1 (Figure 4A) and settled within a few angstroms of the lipid bilayer. The proteins maintained their trimer tertiary structures, and the overall structure of the lipid bilayer was not perturbed by the approach and association of either the wild-type or mutant HR1 domain. We thus took statistics over the last 2 μs of each simulation run to quantify the interaction between the lipid bilayer and the HR1 domain. The wild-type HR1 was consistently located 2-3 Å closer to the bilayer than the 5Q mutant site (Figure 4B), which is consistent with the diminished electrostatic interactions between the 5Q mutant and the charged bilayer. On average, there were 11 PtdIns(4,5)P_2_ molecules within 11 Å of the wild-type HR1 and only 5 PtdIns(4,5)P_2_ molecules within 11 Å of the 5Q mutant (Figure 4C), suggesting that the wild-type Cavin1 HR1 interacts strongly with PtdIns(4,5)P_2_ and concentrates it far above the bulk concentration (of 5% in this case), and the association of PtdIns(4,5)P_2_ with the 5Q mutant is correspondingly decreased. PtdSer association was similar between the wild-type and the 5Q mutant. There are more POPC molecules within 11 Å of the wild-type HR1 than the 5Q mutant, possibly attributable to the larger overall distance of the 5Q mutant from the lipid bilayer. When the lipid interaction cutoff was expanded to 13 Å, both the total number of lipids and the number of POPC molecules interacting with the 5Q mutant were similar to the wild-type, but the number of PtdIns(4,5)P_2_ molecules near the 5Q mutant (6 on average) was still significantly lower than the wild-type. As such, CG-MD modelling supports the view that the cavin1 HR1 domain has a PtdIns(4,5)P_2_-specific interaction with membranes, and this specific interaction is significantly diminished in the 5Q mutant.

We next tested the role of a UC1 domain of cavin1 that has been shown *in vitro* to bind PtdSer ^34^. We generated two cavin UC1 domain mutants: deletion of the entire UC1 domain (deltaUC1) and replacing lysines and arginine residues within the UC1 domain by aspartic acid and glutamate (6KR) (Figure 1A). CAV1 was co-expressed with either cavin1 UC1 mutant in MCF7 cells for another set of lipid mapping analysis. The deltaUC1 mutation completely ablated co-clustering of with PtdSer, increased co-clustering with PA and PtdIns(3,4,5)P_3_, without affecting PtdIns(4,5)P_2_ and cholesterol (Figure 2A,B). Cavin1-6KR mutation also decreased PtdSer co-clustering, but with less impact on other lipids than complete deletion of the UC1 domain. Interestingly substitution of the acidic residues in the UC1 domain also caused a loss of cholesterol co-clustering (Figure 2A,B). These results suggest that the specific lipid environment of caveolae is generated synergistically by both caveolins and by cavins. *In vitro* studies showing changes in PtdIns(4,5)P_2_ or PtdSer binding *in vitro* translate into remarkable changes in lipid association in cells. Moreover, loss of binding to these specific lipids has indirect effects on cholesterol association with CAV1/cavin1 domains to alter the entire lipid environment showing a cooperativity in lipid recruitment to caveolae.

## Association of caveolin and cavin1 requires distinct PtdSer species with unique acyl chain composition

In view of the PtdSer-binding specificity via the cavin UC1 domain that is required for caveola stability and the role of PtdSer in caveola formation ^35^, we next analysed whether PtdSer levels determined CAV1/cavin1 association. PSA3 cells generate less endogenous PtdSer when grown in medium containing dialyzed fetal bovine serum (DFBS), but PtdSer can be restored to near control levels by 10 μM ethanolamine (Etn) supplementation ^16-20,36^. PSA3 cells cultured under different conditions to modulate their PS content were transfected with CAV1 and cavin1 for a bivariate co-localization analysis. Negligible CAV1/cavin1 association was observed in PSA3 cells depleted of PS (DFBS, Figure 5). In contrast, cells grown in Etn with near-native levels of PtdSer showed highly significant association of CAV1 and cavin1. Under the endogenous PS depletion condition, we next added synthetic PtdSer species to PSA3 cells to enrich a specific PtdSer species (Figure 5A). Addback of fully saturated DSPS (di18:0) and mono-unsaturated DOPS (di18:1) were highly effective in driving CAV1 /cavin1 association. In contrast, the mixed chain POPS (16:0/18:1) was ineffective (Figure 5B). These results show that the association of caveolins and cavins is dependent on specific PtdSer species. DOPS, DSPS favor highly curved PM whereas POPS prefers flatter PM ^20^, suggesting that CAV1/cavin1 association may depend on the physical characteristics of PtdSer acyl chains.

## Discussion

This study provides the first quantitative picture of the lipid environment associated with the cytoplasmic face of caveolae *in situ* (for schematic summary see Figure 6). CAV1 and cavin1 in this model system generate caveolae significantly enriched in PtdIns(4,5)P_2,_ PtdSer and cholesterol. This environment is quantitatively distinct from that seen with expression of CAV1 alone (PtdSer, cholesterol, PtdIns(3,4,5)P_3,_ lower PtdIns(4,5)P_2_) or cavin1 alone (PtdIns(3,4,5)P_3,_ PtdIns(4,5)P_2_, PA). Through a combination of mutational and co-expression studies we can start to further define the mechanisms and molecular determinants involved in generating these distinct lipid profiles. The caveolin scaffolding domain has been extensively characterized as an inhibitor of signal transduction pathways *in vitro* and *in vivo* ^24,26^ but the mechanisms involved remain controversial ^37,38^. The isolated domain has lipid binding activity ^29 28^. In the full length protein, the CSD has been suggested to be at least partially buried within the membrane of caveolae ^39^. In mammalian cells the CSD region of CAV1 is accessible to antibodies within the Golgi complex but not in caveolae unless cholesterol is depleted from the plasma membrane ^40^. We now show that the CSD itself has the ability to co-cluster with cholesterol and PtdIns(4,5)P_2_ when expressed in isolation. Moreover, expression of the CSD has a striking effect on the organization of the PM lipids, increasing the nanoclustering and surface levels of cholesterol and PtdIns(4,5)P_2_ while significantly decreasing the surface PtdSer levels. These distinct lipid sorting capabilities may impact numerous signalling pathways.

We also show that two distinct lipid binding sites in cavin1 contribute to the selective lipid sorting of caveolae. The widespread changes in the co-clustered lipids induced by changing these sites, that confer specific lipid binding activity *in vitro*, emphasizes that the lipid composition of the domain is not generated by single binding interactions but cooperative interactions between multiple lipid-binding interfaces and membrane biophysical properties. This is also validated in our MD siomulations. These findings are consistent with studies on other lipid binding proteins, such as the BAR domain proten Bin1 and other PI-interacting proteins ^41^ showing that, rather than 1:1 lipid interactions, a PtdIns(4,5)P_2_-binding domain of higher stoichiometry is generated by localised electrostatic interactions with the protein. This generates a PtdIns(4,5)P_2_ domain with higher capacity to bind other PI-binding proteins (in the above example, dynamin2). A similar principle can apply to caveolae; the high concentration of CAV1 and cavin1, both of which form oligomers, interact with lipids through multiple interaction interfaces in a cooperative fashion to generate a unique lipid profile. Loss of just one of these interfaces, in the case of PtdIns(4,5)P_2_ binding site, can change the entire complement of associated lipids. Moreover, the role of CAV1/cavin1 complex in generating the lipid domain is not simply through increased association of specific lipids; for example, comparison of PtdIns(3,4,5)P_3_ co-clustering across the different combinations of expressed caveolar proteins shows that only with WT CAV1/cavin1 is there a complete exclusion (negligible co-clustering with CAV1) of PtdIns(3,4,5)P_3_ (Figure 2, Figure 6). Here we have focused on the lipid domain generated in the cytoplasmic leaflet of caveolae by cavins and caveolin. However, we suggest that the well-documented transbilayer coupling between lipids such as PtdSer and extracellular leaflet lipids ^42^ can contribute to the generation of the unique glycosphingolipid composition of caveolae.

These findings point to a highly cooperative process of caveola formation involving caveolin, cavin, and lipids. This is further supported by analysis of the lipid requirements for CAV1/cavin association, focusing on PtdSer. We now show that CAV1/cavin1 association is dependent on specific PtdSer species. Protein-protein interactions can contribute to this association, but in this model system this is not sufficient to generate the CAV1/cavin1 domain required for caveola formation. We envisage a coincidence detection mechanism in which multiple low affinity interactions, protein-lipid and lipid-lipid, contribute to formation of the caveolar domain. Analogous to BAR domain association with PtdIns(4,5)P_2_, the arrangement of the membrane lipids embedded in the bilayer and the lipid-binding interfaces of the oligomeric protein complexes may be crucial. Only fully saturated DSPS (di18:0) and mono-unsaturated DOPS (di18:1) but not mixed chain POPS (16:0/18:1) were compatible with CAV1/cavin1 association implicating the lipid’s fatty acyl chains in facilitating these interactions. As different PS species vary in their response to changes in PM curvature ^20^ we speculate that lipid packing geometry is also crucial for the caveolar architecture required to facilitate the CAV1/cavin1-lipid interactions necessary to generate caveolae.

These results not only have implications for understanding the formation of caveolae but also for their function. Flattening of caveolae in response to membrane tension changes ^43^ or other stresses ^21^ has emerged as a crucial aspect of caveolar biology. Our results suggest that the caveola domain is stabilized by multiple synergistic low affinity interactions, dependent on specific lipids, rather than stable protein-protein interactions and can therefore be considered a metastable domain poised to disassemble. Loss of caveolae through genetic ablation or treatments that cause acute flattening induce changes in the nanoscale lipid organization of the PM ^44^. Key lysine residues that form the PtdIns binding site of cavin1 become ubiquitinated upon caveola disassembly, suggesting their decreased interaction with PtdIns(4,5)P_2_ and changing the local lipid environment as cavins dissociate into the cytosol. By comparing the properties of CAV1 expressed alone with CAV1/cavin1 co-expressed together, we show that non-caveolar CAV1, recently termed the CAV1 scaffold ^23^, diffuses more rapidly in the absence of cavin1, as monitored by single molecule tracking, and has a distinct complement of associated lipids. CAV1 domains are enriched in PI(3,4,5)P_3_ and relatively depleted of PI(4,5)P_2_ and PA (Figure 5B) as compared to the CAV1/cavin1 (caveolar) domain. Caveolae disassembly may, thus, released particular lipid types into the bulk membrane and influence the properties of the bulk membrane, which may contribute to the modulation of Ras signalling ^44^ or actin organization at the junctional caveolae enriched with PI(4,5)P_2_ (Teo et al, BioArchives). In view of the high concentration of cavin proteins associated with each caveola (estimated as 50-80; ^22^) and the proposed number of cavin-associated PI(4,5)P_2_ molecules per cavin trimer (Figure 4), as suggested from our coarse grain simulations, disassembly of caveolae could change the accessibility of a considerable pool of PI lipids. Defining the lipid components that associate with caveolae and the mechanisms that dictate their association with, and dissociation from, caveolae will be crucial to understanding how caveolae function but will also provide general insights into the interactions between protein complexes and the hundreds of membrane lipid species required for cellular function.

## ACKNOWLEDGEMENTS

This work was supported by the National Health and Medical Research Council of Australia (grants APP1140064 and APP1150083 and fellowship APP1156489 to RGP and GNT1120381 GNT1155794 (fellowship to FAM)). BMC is supported by an NHMRC Senior Research Fellowship (APP1136021). NA is supported by an NHMRC grant (APP1102730). RGP is supported by the Australian Research Council (ARC) Centre of Excellence in Convergent Bio-Nano Science and Technology. YZ and JFH are supported by the National Institue of Health (USA) grant R01 GM124233. Computational resources for the CG-MD simulations were provided by the Research Computing Centre at the University of Queensland.

## Supplementary Figures

**Figure S1.**
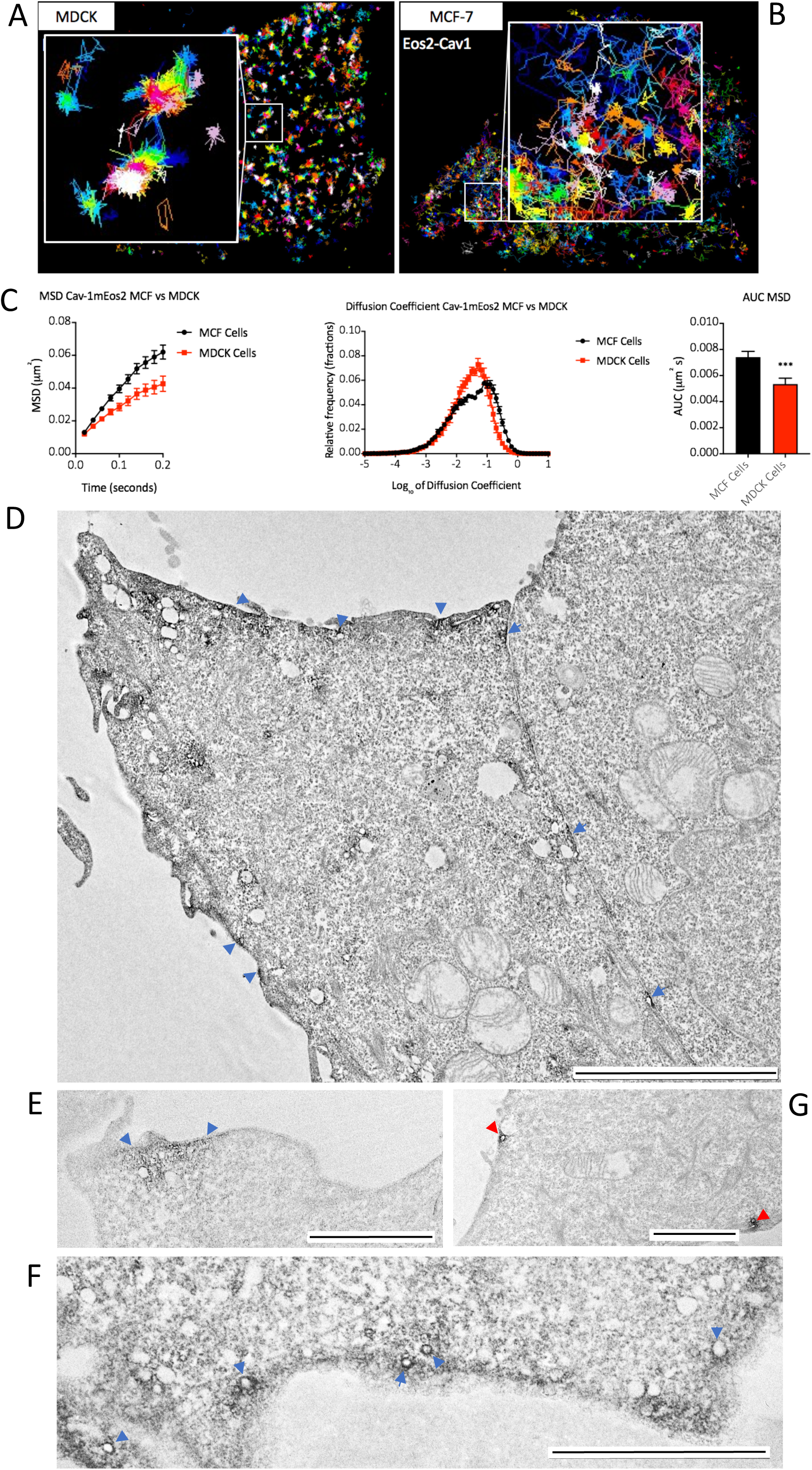
Characterization of the non-caveolar pool of CAV1. Single molecule tracking of CAV1-mEos2 in A. MDCK cells and B. MCF7 cells. C; quantitative comparison of diffusion of CAV1-mEos2 in MDCK vs MCF7 cells showing changes in i.) MSD (µm^2^), ii.) area under the MSD curve and iii.) relative frequency distribution of diffusion coefficient. D, E, F; YFP-CAV1/APEX-nanobody expressed in MCF7 cells. Blue arrowheads indicate plasma membrane areas with high concentration of APEX staining. A range of structures are labeled including flat areas of the plasma membrane (D), clusters of vesicles (E, F) and vesicular profiles or various diameters (F); G) YFP-CAV1/Apex-nanobody co-expressed with cavin1 in MCF7 cells; vesicular profiles characteristic of caveolae are positive for the APEX reaction product (red arrowheads) with negligible labeling elsewhere on the plasma membrane. Bars D, 5µm; E,F,G;2µm.

**Figure S2.**
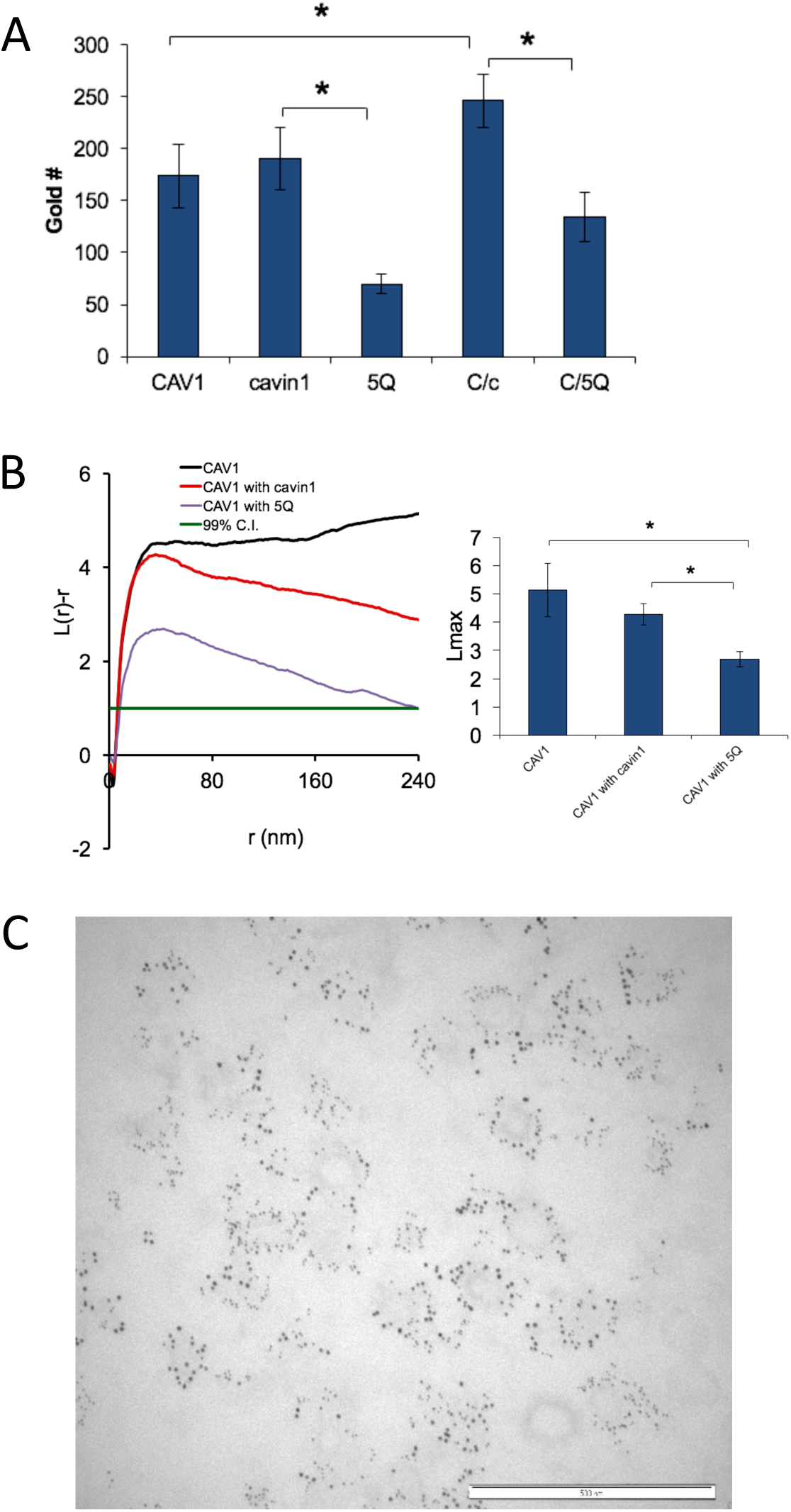
Characterization of the MCF7 system. A. Quantitative analysis of gold labeling density for CAV1, cavin1 and the cavin1.5Q mutant (5Q) or for CAV1 co-expressed with wild type cavin1 (C/c) or cavin1.5Q (C/5Q). B. Univariate analysis of clustering of the expressed constructs. C; unlabeled version of Figure 1D.

## Materials and Methods

### Cell culture and transfection

MCF7 and MDCK cells were obtained from ATCC and regularly mycoplasma tested. MCF7 cells were checked by Western blotting for lack of CAV1 (note that unlike the line used here some strains of MCF7 cells do express low levels of CAV1). MDCK cells were grown in DMEM/F-12 (Life Technologies, Carlsbad, CA, USA) supplemented with 10% FBS and 2 mM L-glutamine. MCF-7 cells were grown in DMEM supplemented with 10% FBS and 2 mM L-glutamine. Caveolin-1 and cavin1 were cloned as described in ^***7***^. The CSD construct ^45^ and cavin constructs ^30 34^ were described previously. Tagged constructs were transfected using Lipofectamine 2000 reagent (Life Technologies, CA, USA) following the manufacturer’s instructions using a 1:3 ratio of DNA:Lipofectamine. 3All transient transfections were performed with Lipofectamine 2000 (Life Technologies) as per the manufacturer’s instruction. PSA3 cells were a generous gift from Dr. Hiroyuki Arai at the University of Tokyo, Japan. PSA3 cells were cultured in DMEM containing 10% dialyzed FBS (DFBS) without/with 10 μM ethanolamine (Etn) for 72 hours before experiments.

#### Acute exogenous PtdSer addback

Synthetic PtdSer species were purchased from Avanti Polar Lipids, Inc. (Alabaster, AL, USA), dissolved in chloroform and kept in nitrogen at −20 °C. Appropriate amount of PtdSer/chloroform solution was transferred to a glass vial (for a final working concentration of 10 μM) using a Hamilton syringe. Chloroform was evaporated by purging with nitrogen and kept under vacuum overnight in the dark to eliminate residual chloroform. The dried lipid film was rehydrated in DMEM containing 10% DFBS and sonicated for 20 minutes in a bath sonicator. The final medium containing PtdSer suspension was added to PSA3 cells for one hour before harvesting.

### Electron Microscopy

Standard transmission EM using ruthenium red and immunoelectron microscopy on frozen sections using antibodies to CAV1 followed by 10nm protein A-gold were performed as described previously ^7^. Sonication of MCF7 cells to generate plasma membrane lawns was performed exactly as described ^22^. Briefly cells were plated on poly-L-lysine coated 35mm plastic dishes and then a probe sonicator was used to generate basal PM sheets adhered to the substratum. Cells were then fixed and labeled before embedding in resin and sectioning close to the substratum. Localization of YFP-CAV1 using a nanobody-based APEX2 approach was as described previously ^46^.

### Quantitative spatial analysis

#### Univariate analysis

The univariate analysis calculates the spatial distribution of a single population of gold nanoparticles within a select PM area ^16,19^. The GFP-tagged proteins/peptides of interest on intact PM sheets were attached to copper EM grids, fixed with 4% paraformaldehyde (PFA) and 0.1% gluaraldehyde, and immunolabeled with 4.5nm gold conjugated to anti-GFP antibody and negative-stained with uranyl acetate. Gold nanoparticles were imaged with TEM at 100,000x magnification. ImageJ was then used to assign x, y coordinates for each gold particle. The spatial distribution of gold particles within a selected 1μm^2^ area was calculated using the Ripley’s K-function, which tests a null hypothesis that all points in the analyzed area are randomly distributed:

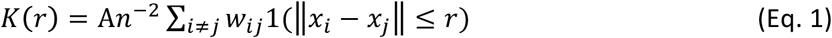

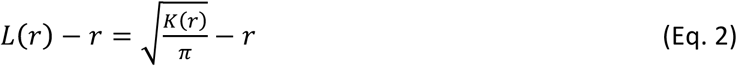

where *K(r*) describes the univariate K-function for the total number of gold particles, denoted as *n*, in a selected area of *A*; *r* denotes the distance between 1 and 240 nm with an increment of 1 nm. ‖ ^**·**^ ‖ characterizes Euclidean distance, with 1(^**·**^) as the indicator function. We define: 1(^**·**^) of = 1 if ‖*x*_*i*_-*x*_*j*_‖ ≤ r and 1(^**·**^) = 0 if ‖*x*_*i*_-*x*_*j*_‖ > r. We incorporate a parameter of *w*_*ij*_^-1^ to achieve an unbiased edge correction. For a circle with *x*_*i*_ at the center and ‖*x*_*i*_-*x*_*j*_‖ as the radius, *w*_*ij*_^-1^ defines the proportion of the circumference of the circle. *L*(*r*) – *r* is a linear transformation of *K*(*r*), and is normalized against the 99% confidence interval (99% C.I.) calculated from Monte Carlo simulations. A *L*(*r*) - *r* value of 0 describes a complete random pattern of particles distribution, while a *L*(*r*) - *r* value above the 99% C.I. of 1 illustrates a statistically meaning clustering pattern. For each condition, at least 15 PM sheets were imaged and analyzed. Statistical significance between differet conditions was evaluated by comparing our calculated distribution patterns against 1000 bootstrap samples in bootstrap tests ^16,19^.

#### Bivariate co-clustering analysis

Co-clustering between two different sized gold particles is calculated using the bivariate K-function co-clustering analysis ^16,19^. The GFP- and RFP-tagged proteins on intact PM sheets attached to EM grids were co-immunolabeled with 2nm gold conjugated to anti-RFP antibody and 6nm gold linked to anti-GFP antibody, respectively. As above, the x, y coordinates of gold particles were assigned using ImageJ. Bivariate K-function analysis then calculated the co-localization between 6 nm and 2 nm gold populations. The null hypothesis of this bivariate analysis is that the two gold populations spatially separate from each other. (Eqs. 3-6):

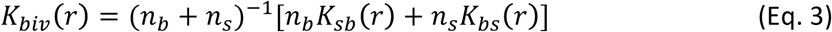

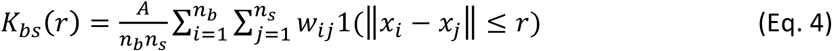

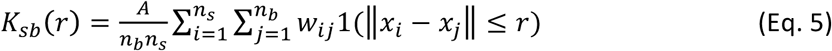

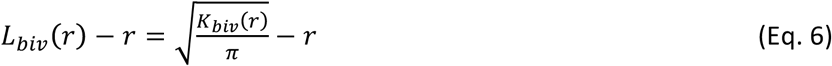

where *K*_*biv*_(*r*) denotes the bivariate estimator containing 2 distinct bivariate K-functions: *K*_*bs*_(*r*) calculates the distribution of all the 6 nm gold particles (*b* = big gold) with respect to each 2 nm-small gold (*s* = small gold); and *K*_*sb*_(*r*) calculates the distribution of all the 2 nm gold with respect to each 6 nm gold. The total number of 6 nm big gold is termed as n_b_, while the total number of 2nm small gold n_s_. Other notations for equations 3-6 still follow the same description as in Eqs.1 and 2. *L*_*biv*_(*r*)-*r* is a linear transformation of *K*_*biv*_(*r*), and is further normalized against the 95% C.I. An *L*_*biv*_(*r*)-*r* value of 0 indicates lateral separation between 6nm/2nm gold particles. On the other hand, *L*_*biv*_(*r*)-*r* values above the 95% C.I. of 1 indicate statistically meaningful co-localization. To better summarize the co-localization data, we integrate the *L*_*biv*_(*r*)-*r* curves within a fixed range 10 < *r* < 110 nm, and termed the parameter as bivariate *L*_*biv*_(*r*)-*r* integrated (or LBI):

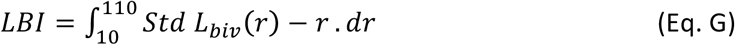

For each test, at least 15 PM sheets were imaged and analysed. The same bootstrap tests used above in the univariate analysis are also used here to evaluated the statistical significance between tests ^16,19^.

### Single molecule tracking and quantitation

Single particle tracking photoactivated localization microscopy (sptPALM) of MCF and MDCK cells transfected with Cav1-mEos2 was carried out on the Roper Scientific TIRF microscope equipped with an iLas^2^ double laser illuminator (Roper technologies), a CFI Apo TRIF 100× (1.49-NA) objective (Nikon) and an Evolve512 delta EMCCD camera (Photometrics). Images were acquired using Metamorph software (version 7.78; Molecular Devices) at 50 Hz and 16000 frames were acquired per cell. A 405 nm laser was used to photoconvert mEos2, with simultaneous 561 nm exposure to excite the photoconverted mEos2. For stochastic photoconversion of mEos2 molecules, low amount (3-5%) of 405 nm laser and 75-80% of 561 nm laser was used. Data analysis was carried out as previously described ^47,48^ using PALM-Tracer, a plugin in Metamorph software (Molecular Devices).

### Molecular simulations

The interaction between Cavin1-HR1 and lipid bilayers was modelled using the MARTINI 2.2 force field ^31-33^. The RCSB PDB structure ID 4QKV was used as the starting point for the Cavin1-HR1 structure, with the 5Q point mutations performed using the Mutate Residue tool in VMD ^49^. Both Cavin1-HR1 and Cavin1-HR1-5Q atomistic structures were then coarse-grained using the martinize.py script ^31^ with default parameters. Using the insane.py script ^50^, the resulting coarse-grained protein structures were horizontally aligned and placed 4 nm away from a coarse-grained 80:15:5 POPC:POPS:PI(4,5)P2 lipid bilayer with 840 lipids per leaflet. The simulation unit cell was cubic with initial size 24 × 24 × 20 nm^3^.

All simulations were performed in GROMACS 2019.3 ^51^ using standard MARTINI 2.2 parameters ^52^ [with van der Waals Lennard-Jones potentials shifted to zero at a cutoff of 1.1 nm and long-range Coulomb interactions treated with a reaction-field with a cutoff of 1.1 nm and dielectric constant ε_r_ = 15]. Each of four runs began with 10000 steps of energy minimization, followed by 0.5 ns of NVT equilibration and 1 ns of NPT equilibration, using a time step of 20 fs, V-rescale thermostat at 300K with time constant 1 ps and (for the second equilibration) Berendsen semi-isotropic barostat at 1 bar with time constant 4 ps and compressibility 4.5 × 10^−5^ bar^-1^. The production run was then carried out for 3 microseconds, using a time step of 30 fs, V-rescale thermostat at 300K with time constant 1 ps and Parrinello-Rahman semi-isotropic barostat at 1 bar with time constant 12 ps and compressibility 3.0 × 10^-4^ bar^-1^.

Protein-bilayer distances and lipid occupancies were calculated using GROMACS tools gmx mindist and gmx select respectively. Distances were measured as the minimum between any protein particle and any lipid particle, while occupancies were measured by counting all lipids with at least one particle within the occupancy cutoff of any protein particle.

